# Separating the effects of sex hormones and sex chromosomes on behavior in the African pygmy mouse *Mus minutoides*, a species with XY female sex reversal

**DOI:** 10.1101/2022.07.11.499546

**Authors:** Frederic Veyrunes, Julie Perez, Louise Heitzmann, Paul A Saunders, Laurent Givalois

**Author notes:** Corresponding author: Frederic Veyrunes.

## Abstract

In mammals, most sex differences in phenotype are controlled by gonadal hormones, but recent work on transgenic mice have shown that sex chromosomes can have a direct influence on sex-specific behaviors. In this study, we take advantage of the naturally occurring sex reversal in a mouse species, *Mus minutoides*, to investigate for the first time the relationship between sex chromosomes, hormones and behaviors in a wild species. In this model, a feminizing variant of the X chromosome, named X*, produces three types of females with different sex chromosome complements (XX, XX*, and X*Y), associated with alternative behavioral phenotypes, while all males are XY. We thus compared the levels of three major circulating steroid hormones (testosterone, corticosterone and estradiol) in the four sex genotypes to disentangle the influence of sex chromosomes and sex hormones on behavior. First, we did not find any difference in testosterone levels in the three female genotypes, although X*Y females are notoriously more aggressive. Second, in agreement with their lower anxiety-related behaviors, X*Y females and XY males display lower baseline corticosterone concentration than XX and XX* females. Instead of a direct hormonal influence, this result rather suggests that sex chromosomes may have an impact on the baseline corticosterone level, which in turn may influence behaviors. Third, estradiol concentrations do not explain the enhanced reproductive performance and maternal care behavior of the X*Y females compared to the XX and XX* females. Overall, this study highlights that most of the behaviors varying along with sex chromosome complement of this species are driven by genetic factors rather than steroid hormone concentrations.

## INTRODUCTION

The developmental processes leading from a single genome to two sexually dimorphic phenotypes are remarkably complex and remain poorly understood (Mank, 2017; Naqvi et al., 2019). In mammals, sex development can be divided into two main components: sex determination and sex differentiation. Sex determination is the process by which the bipotential embryonic gonads develop into either testes or ovaries, which depends exclusively on the sex chromosomes; sex differentiation involves the subsequent events that ultimately produce either the male or female phenotype, driven mostly by gonadal steroid hormones (review in Arnold, 2012). However, in the last two decades, several studies on genetically manipulated mice have challenged this view, highlighting the direct influence of the sex chromosomes on sex differences in behavior and brain, such as some aspects of aggressiveness, parental behavior, food consumption, social interactions, anxiety, or brain structures and functions (e.g. de Vries et al., 2002; Gatewood et al., 2006; Arnold, 2012, 2019; Lentini et al., 2013; Corre et al., 2016). Although mammalian XX/XY sex determination is extremely conserved, a few species escape this convention and present unusual sex determination systems (Parma et al., 2016; Saunders & Veyrunes, 2021). The African pygmy mouse, *Mus minutoides* is the latest addition to this short list of odd mammals, owing to the presence of a third sex chromosome named X* (Veyrunes et al., 2010). This is a mutant copy of the X chromosome that differs cytogenetically from the ancestral X and is feminizing. Therefore, three female genotypes are found: XX, XX* and X*Y, while all males are XY. Previous works showed that sex-reversed females (X*Y) may represent up to 75% of the females in some wild populations (Veyrunes et al., 2010). They are fully fertile with typical ovaries without apparent signs of testicular tissue (Rahmoun et al., 2014), they have a higher lifetime reproductive success than the XX and XX* females, including larger litter size despite the cost of losing lethal YY embryos (Saunders et al., 2014), and they also show enhanced maternal care behaviors compared to other females (Heitzmann et al. 2022). On the other hand, they display some masculinized behaviors, such as enhanced aggressiveness, lower anxiogenic response (Saunders et al., 2016) and higher bite-force performance (Ginot et al., 2017). Therefore, the X* effect goes well beyond sex reversal, and is at the origin of a complex X*Y phenotype with opposite results on the two components of sex development (i.e., sex determination and sex differentiation): gonad feminization and partial masculinization of behaviors. Phenotypic sexes of the X*Y gonads and brain are thus partially dissociated, a unique feature in a wild mammal species. Moreover, gonadal sex and chromosomal sex also are partially uncoupled (XX *vs*. X*Y females, and XY males *vs*. X*Y females), offering a promising model to investigate the influence of sex chromosome complements versus steroid hormones on sexual dimorphism in brain and behavioral traits. To date, the behavioral differences observed in the three *M. minutoides* female genotypes have been mostly attributed to genetic effects (Saunders et al., 2016; Ginot et al., 2017) on basis that X*Y females are characterized by complete sex reversal with typical ovarian anatomy (no ovotestes) and normal anogenital distance. This suggests a low level of circulating androgens (Rahmoun et al., 2014), but the question remains open. In this study, we thus compared the levels of three circulating steroid hormones (testosterone, corticosterone and estradiol) in the four sex genotypes (XY males, and XX, XX* and X*Y females) to disentangle the influence of sex-linked genes and steroid hormones on the masculinized behaviors observed in X*Y females (Saunders et al., 2016; Ginot et al., 2017).

## MATERIALS AND METHODS

### Animals

Mice were bred in our own breeding colony (CECEMA, University of Montpellier), established from wild-derived animals (for further details see Saunders et al., 2014). Females were housed in same-sex groups of 3-4 individuals per cage, and males in individual cages (to prevent agonistic behaviors). They were provided with *ad libitum* food and water, and the light regime was set to 15:9h (light:dark). The genotype of females was determined by PCR amplification of the Y-specific *Sry* gene from tail biopsies or culture of fibroblasts from skin biopsies. The animal age varied between 8 and 10 months, and all blood/urine samples were collected at the same time of the day (± 2 h). Experimental procedures were in accordance with the European guidelines and were approved by the French Ethical Committee on animal care and use (No. CEEA-LR-12170).

### Hormone assays

As the African pygmy mouse is one of the smallest mammal species in the world (Boel et al., 2020), the amount of blood that could be collected from each individual was limited. This prevented testing all samples in duplicate and measuring the three steroid hormones in the same animal (one mouse for each hormone). Moreover, for a limited number of animals, reduced volumes of plasma, serum or urine were collected and had to be used diluted with kit’s sample diluents or SAMPLEDIL (ref: KLZZ731). Nevertheless, all samples were within the range of robust minimal assay sensitivity. The numbers of animals used for the different measurements are listed in Table 1.

**Table 1:** Number of mice involved in each hormone assay.

| Sex | Female |  |  | Male |
| --- | --- | --- | --- | --- |
| Genotype | XX | XX* | X*Y | XY |
| Testosterone | 10 | 10 | 10 | 10 |
| Corticosterone (baseline) | 8 | 10 | 10 | 13 |
| Corticosterone (after stress) | 8 | 12 | 13 | 11 |
| Estradiol (diestrus) | 12 | 11 | 13 | NA |
| Estradiol (estrus) | 13 | 13 | 12 | NA |

#### Testosterone

animals were euthanized by decapitation and trunk blood collected in Eppendorf tubes. After 30 min at room temperature, blood samples were centrifuged and serum was collected and stored at -20 °C. Testosterone concentrations were measured using an ELISA kit (IBL International ref: RE52151) following the manufacturer’s instructions. To fall within the sensitivity range of the assay, serum from males were tested at two dilutions (1:5 and 1:10), and the hormone concentration was calculated by the mean of values obtained at each dilution. This also allowed to obtained pseudo-duplicates that confirmed an excellent dilution pattern, and a very good assay precision with a low intra-assay coefficient of variability.

#### Corticosterone

plasma corticosterone concentrations were measured at baseline (no stress) and after stress induction. For baseline corticosterone, mice were taken from the cage and rapidly decapitated to collect trunk blood. For the post-stress corticosterone measurement, mice were placed in a glass beaker (diameter 12.5 cm), filled with 1.4L of water (22 ± 1 °C) and forced to swim for 10 min, then animals were rapidly decapitated. In both cases, blood samples were collected in EDTA-coated Eppendorf tubes, centrifuged at 4 °C, and plasma was collected and stored at -20°C. Corticosterone concentrations were measured with a conventional ELISA kit (ENZO Life Sciences; ref: ADI-900-097) following the manufacturer’s instructions, as previously reported (Canet et al., 2020).

#### 17ß-Estradiol

the inconclusive results obtained with serum (most concentrations below the standard curve) led to its quantification in urine samples that have shown to give interpretable results in mice (e.g. deCantazaro et al., 2004). Urine was collected upon handling mice and pipetted on a clean surface, and stored at -20 °C. As estradiol concentration varies along the estrous cycle (e.g. Wood et al., 2007), this study focused arbitrarily on the diestrus and estrus stages. To determine the estrous cycle phase, just after urine collection, females were held by the tail, and the vagina was flushed twice with 10 µl of sterile water using a pipette tip. The vaginal smear was placed on a glass slide, stained with Giemsa, and viewed under a light microscope (10x magnification). The diestrus phase was defined by the exclusive presence of leukocytes, and the estrus phase by the presence of large and squamous-type epithelial cells without nucleus. Urinary estradiol concentrations were measured with an ELISA kit (IBL International ref: 30121045) following the manufacturer’s instructions and protocol in deCantazaro et al. (2004).

### Statistical analysis

Final steroid concentrations were calculated from hormone-specific standard curves corrected for an individual’s sample volume. All statistical analysis were performed on R v3.6.3. We used a non-parametric test, Kruskall-Wallis, to test differences in testosterone level between the four genotypes. Post-hoc pairwise comparisons were made using *dunn*.*test* with a Benjamini-Hochberg p-value adjustment to account for multiple testing. Baseline and stress-induced corticosterone concentration were analyzed separately fitting a linear model with individuals’ genotype as the only predictor. We used a log-transformation of corticosterone baseline-level to account for heteroscedasticity. Post-hoc pairwise comparisons were made using *emmeans* package with Tukey adjustments of p-values to account for multiple testing. We also fitted a linear model to investigate estradiol level using ‘genotype’ and ‘estrous cycle phase’ (estrus *vs*. diestrus) as predictors. We included an interaction between both predictors with the assumption that female estradiol level may vary with the estrous cycle and independently between genotypes (i.e., estradiol variation between estrus and diestrus states may differ between genotypes). Post-hoc comparisons were made using *emmeans* package with Tukey adjustments of p-values to account for multiple testing.

## RESULTS AND DISCUSSION

Sex-reversed *M. minutoides* females show a combination of phenotypic traits not seen in concordant individuals of either sex. For some phenotypic traits, X*Y females can be viewed as “super females” as called by Knight (2017), with a greater reproductive output (Saunders et al., 2014) and enhanced maternal care (for example, they retrieve pups faster and with a higher probability; Heitzmann et al. 2022) compared to XX and XX* females. Conversely, for other aspects, they are more similar to males: lower anxiety-related behavior and enhanced aggressiveness (Saunders et al., 2016). That similarity between X*Y females and males extends to whole-organism performance with higher bite force than other females, and even a tendency (though not statistically significant) to bite harder than males (Ginot et al., 2017). In summary, sex reversal changes not only the gonadal sex, but also some behavioral and life history traits, and is at the origin of a new female sexual phenotype (Heitzmann et al., 2022). Therefore, to untangle the role of genetic *versus* hormonal effects on these behavioral differences observed among females, we compared the levels of three major circulating sex steroid hormones in the different genotypes.

Testosterone is ubiquitous in vertebrates and is the key mediator in the expression of many male morphological and behavioral traits (Trainor and Marler, 2002; Giammanco et al., 2005; Preston et al., 2012; McEvoy et al. 2015; and references therein). More particularly, it is thought to be the major sex steroid regulating male aggressive behavior and dominance, in a wide variety of fish, reptile, bird, and mammal species (including humans). For example, high rates of aggression generally correlate with elevated testosterone levels (Giammanco et al., 2005; Yu and Shi, 2009; Cantarero et al., 2015; St Clair Yewers et al., 2017; Denson et al. 2018), and male aggressiveness can be reduced by castration and restored by administration of androgens (Giammanco et al. 2005). In humans, higher testosterone concentrations are observed in perpetrators of violent crimes, in soldiers with antisocial behaviors, in subjects with impulsive behaviors, and in athletes during competition (review in Giammanco et al., 2005). In vertebrates, testosterone is mostly produced by Leydig cells in testes, while smaller quantities are also produced by ovaries, and adrenal glands in both sexes (Nelson, 2010). Although aggressiveness and testosterone may be lower in females, the more limited literature data shows a similar positive relationship between testosterone and aggression, albeit less so (Giammanco et al., 2005; Denson et al., 2018). In the African pygmy mice, we found a high sexual dimorphism of serum testosterone levels as expected (p < 0.001 between males and XX and XX* females; p = 0.002 between males and X*Y females); however, we did not detect any difference among the three female genotypes (Fig. 1). Therefore, the aggressive behavior and stronger bite force (another indicator of aggressive performance) of X*Y females are unlikely to be directly mediated by the circulating testosterone level. This result adds to the increasing evidence that the relationship between aggression and testosterone is not straightforward (e.g. Huyghe et al., 2009; McEvoy et al., 2015; Cantarero et al., 2015; St Clair Yewers et al., 2017), which might be explained by the fact that aggressiveness is highly context-dependent (e.g. social experience or motivation) and includes complex behavioral patterns with different underlying physiological mechanisms and metabolic pathways. This is especially relevant in females: in females more than males, it would be simplistic to reduce hormone-mediated aggressive behavior to testosterone concentration only (review in Denson et al., 2018). In *M. minutoides*, the clear-cut contrast of aggressiveness in controlled experiments between X*Y females and XX and XX* females (in the resident-intruder test, X*Y females show an attack latency at least twice shorter and a number of aggressions at least twice higher than in the other female genotypes; Saunders et al., 2016) may also suggest a direct effect of the sex chromosome complement on the behavior of X*Y females. Specifically, the Y chromosome includes at least one gene, *Sts*, that has been shown to influence aggressiveness in mice (Mortaud et al., 2010). Two other important candidates are X-linked genes, the Androgen Receptor (*AR*) which mediates the actions of testosterone (Davey & Grossman 2016), and the Monoamine oxidase A (*MaoA*) that is well known to influence aggressiveness (Cases et al., 1995). Following the rise of the X* chromosome, differences of expression could have evolved between the X and X* copies.

**Fig. 1:**
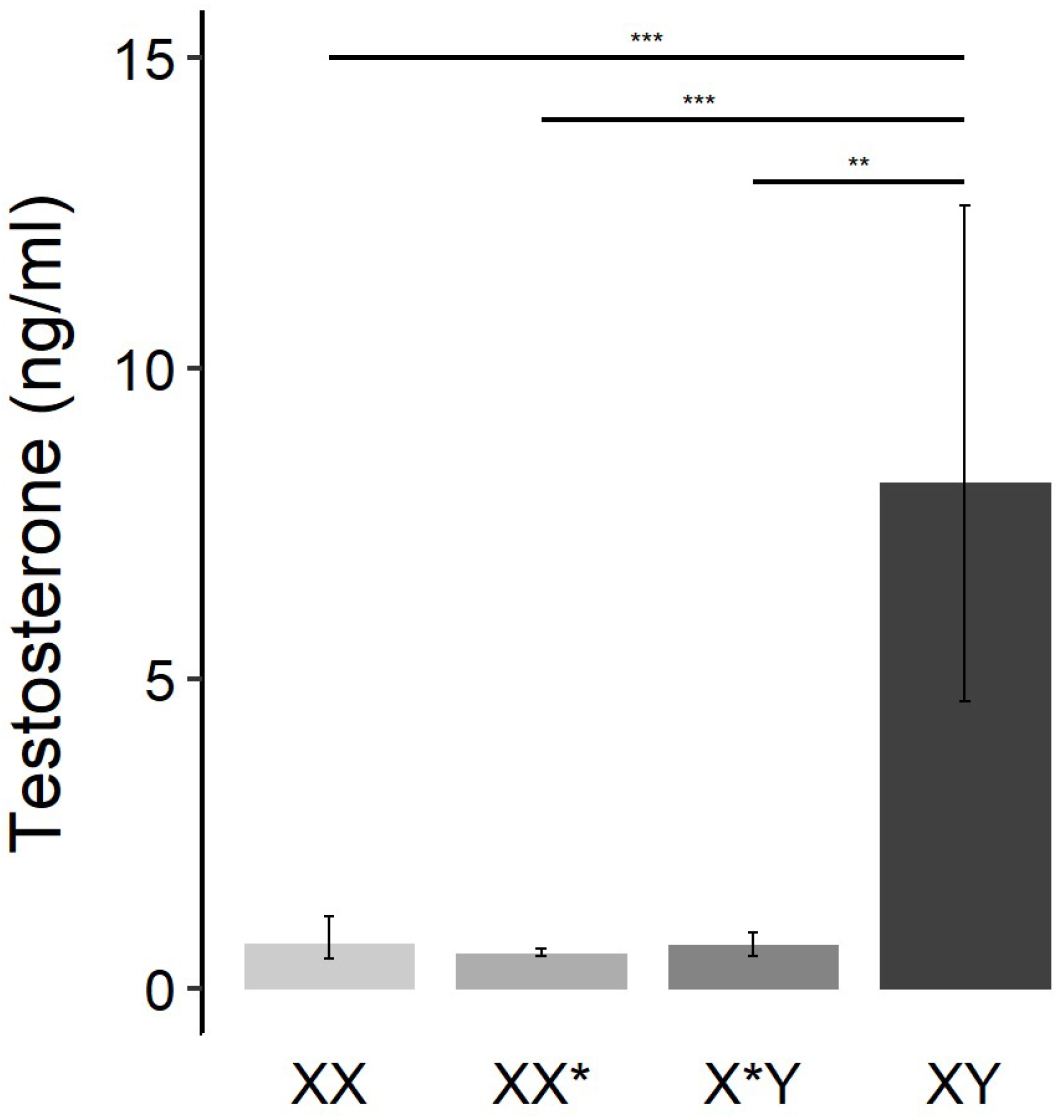
Serum testosterone concentrations of the four sex genotypes, XX, XX* and X*Y females, and XY males. Values are means ± IC 95 %. *** p < 0,001, ** p < 0,01.

Although glucocorticoids (i.e., mainly corticosterone in rodents) may be involved in aggression, they are best known as stress hormones (Romero, 2004; Bonier et al., 2009). Corticosterone secreted by adrenal glands is increasingly employed as a physiological indices of the relative health conditions and as a stress index of individuals or populations (review in Bonier et al., 2009). The interpretation of corticosterone concentrations is complicated by the fact that its secretion can be classified in two types based on the secretion dynamics, the receptors bound, and the physiological and behavioral effects (Romero, 2004; Rensel and Schoech, 2011). The baseline corticosterone is the amount of hormone secreted on a continuous low-level basis, and is involved in the maintenance of energetic balances and homeostasis. Elevated baseline corticosterone levels are assumed to predict a low relative fitness of individuals, associated with multiple deleterious characteristics, such as chronic stress, poor body conditions, reallocation of resources away from reproductive behaviors, infertility (e.g. Romero and Wikelski, 2001; Strasser and Heath, 2013; review in Bonier et al., 2009). The second type of secretion takes place when a stressful stimulus is perceived, stress responses are thus initiated which trigger activation of the hypothalamo-pituitary-adrenal axis with a rapid and intense release of corticosterone. This abrupt increase of circulating corticosterone modulates specific physiological and behavioral traits to help the organism to cope with the stressor, thus promoting short-term survival, and will remain high for a few hours (Wingfield and Sapolsky, 2003; Romero, 2004). Acute elevation of corticosterone has been associated with escape behavior, suppression of territorial behavior and reproductive activity, reduction of parental care and energy reallocation (Wingfield et al., 1998; Sapolsky et al., 2000; Miller et al. 2009). Therefore, differences in baseline or stress-induced corticosterone concentrations among individuals may have completely different physiological and behavioral consequences, and it is essential to consider both. In *M. minutoides*, X*Y females and XY males display lower baseline corticosterone levels than XX and XX* females; concentrations were significantly lower in sex-reversed females than in XX* (p = 0.003) and XX females (p = 0.01), tended to be lower though not significant in males than in XX* (p = 0.054) and XX females (p = 0.14), and were similar between X*Y females and males (p = 0.54), as well as between XX and XX* females (p = 0.99); whereas the stress-induced concentrations were comparable in the four groups (Fig. 2A-B). According to the Glucocorticoid-Fitness hypothesis (Bonier et al., 2009), the lower baseline corticosterone concentration in X*Y females suggests lower anxiety and greater relative fitness. This assumption is congruent with the life-history and behavioral traits measured in captivity: X*Y females display (i) a greater probability of breeding and produce significantly more offspring during their lifetime than XX and XX* females (Saunders et al., 2014), and (ii) lower anxiety-related behavior compared with the other females but similar to males (Saunders et al., 2016). This might suggest that anxiety-related behaviors are influenced by sex hormones only; however, the finding that animals harboring a Y chromosome have a lower baseline corticosterone than XX and XX* females, rather suggests a direct effect of the sex chromosome complements. We show that irrespective of the gonadal sex, the chromosomal sex also may have an impact on steroid concentrations, which in turn may influence behaviors.

**Fig. 2:**
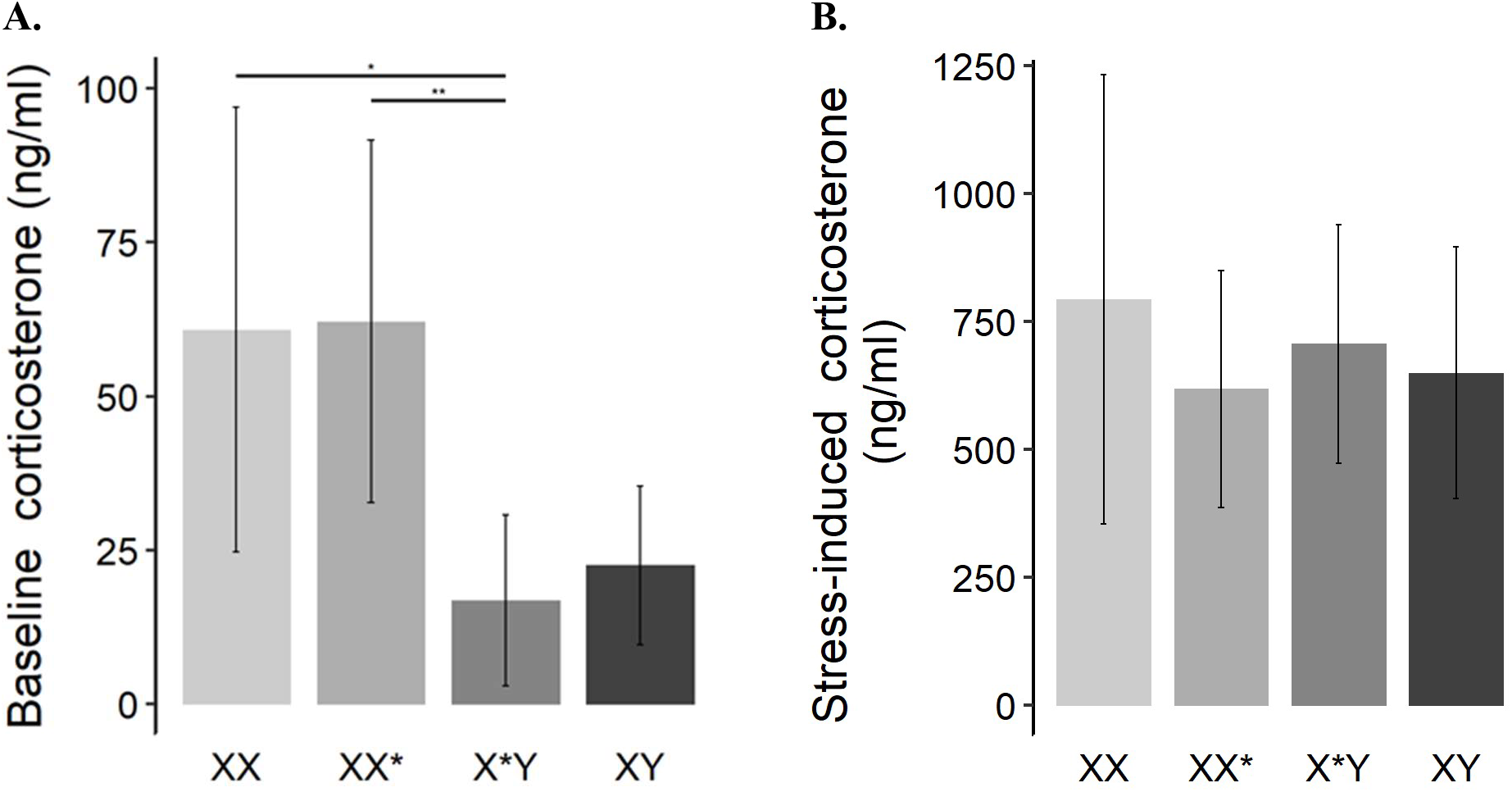
Plasma corticosterone concentrations of the four sex genotypes, XX, XX* and X*Y females, and XY males. (**A**) Baseline and (**B**) stress-induced concentrations. Values are means ± IC 95 %. **p < 0,01, *p < 0,05.

Finally, we investigated estradiol levels in females since the reproductive performances (Saunders et al., 2014) and the maternal care behavior (Heitzmann et al. 2022) of the three female genotypes differ. Estrogens, particularly estradiol, exert powerful influences on the mammalian female reproductive physiology and behavior. For example, estradiol plays critical roles in regulating estrous cycling, preparing the uterus for embryo implantation, inducing sexual receptivity and facilitating lordosis, and on the onset and maintenance of maternal behaviors (reviews in Kelley & Brenowitz, 2002; deCatanzaro, 2015). In *M. minutoides*, urinary estradiol levels were comparable among female genotypes at diestrus, and marginally lower in XX females during estrus (p = 0.053 and p = 0.11 in comparison to X*Y and XX*, respectively) (Fig. 3). Our data also show a trend for higher estradiol levels at estrus than diestrus in X*Y (p = 0.048) and XX* females (p = 0.08), as reported in the house mouse (e.g. Wood et al., 2007), but not in XX females (p = 0.92) (Fig. 3). In our laboratory colony, XX females show lower breeding performance and lower pup retrieval behavior than X*Y females (Saunders et al., 2014; Heitzmann et al. 2022), which fits with the trend in estradiol level differences observed. However, the finding that XX* have comparable reproductive traits and maternal care behaviors to XX females, but the same estradiol concentration as X*Y females rules out any causal link. Conversely, these results strongly suggest an advantageous feature of the X*Y complement, and it is tempting to attribute it to the X* chromosome that evolved towards a female uniparental transmission and may have accumulated female-beneficial genes. But this hypothesis does not explain why XX* females, which also harbor the X* chromosome and retain random inactivation of the X and X* chromosomes (Veyrunes and Perez, 2018), do not present a more intermediate phenotype. Other hypotheses are discussed by Saunders et al (2014), particularly concerning the X chromosome copy number (one in X*Y *vs*. two in XX and XX*) that has been shown to influence the expression of hundreds of autosomal genes in the house mouse (Wijchers et al., 2010). Finally, the low level of estradiol concentration and the absence of detectable variations between estrus and diestrus in XX females are noteworthy, and might be related to their apparent smaller ovaries and fewer follicles compared to XX* and X*Y females (Rahmoun et al., 2014); this result requires further investigations.

**Fig. 3:**
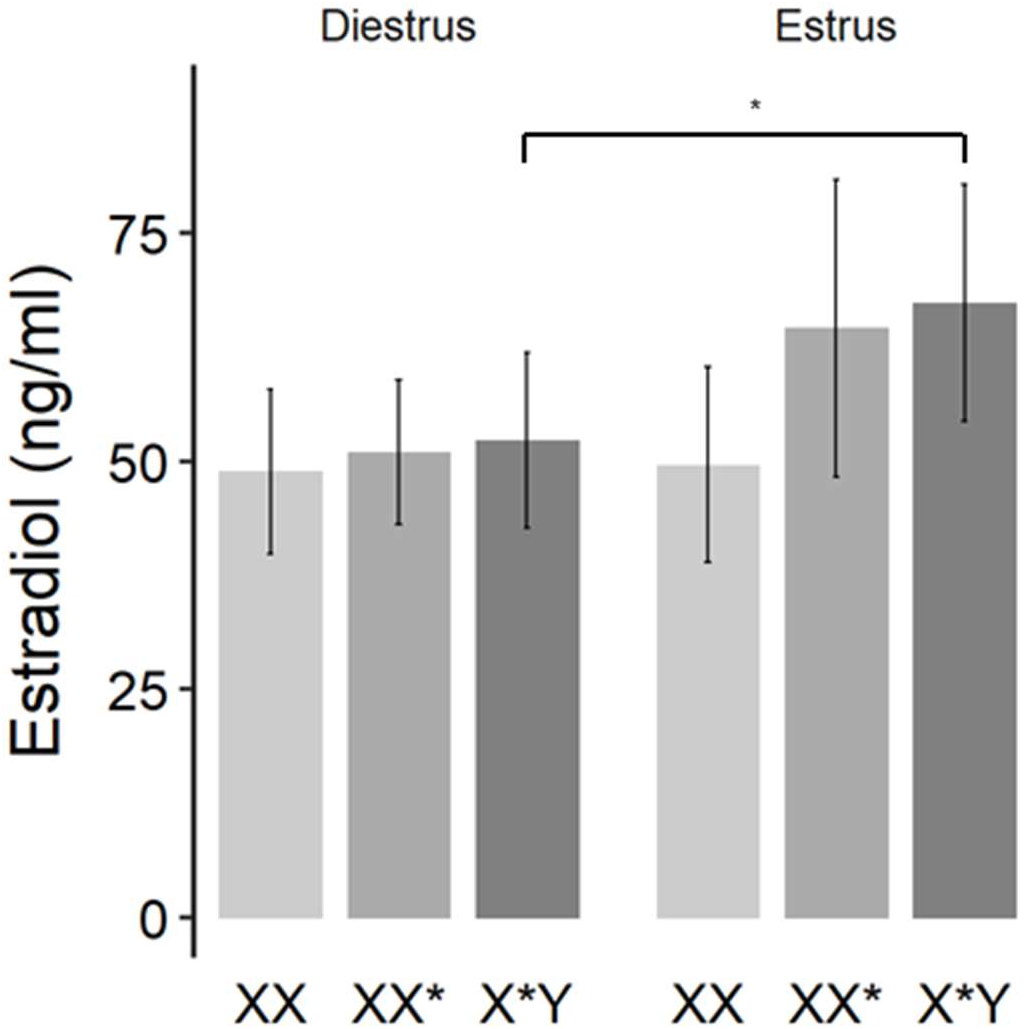
Urinary estradiol concentrations of the three female genotypes, XX, XX* and X*Y, at diestrus and estrus stages. Values are means ± IC 95 %. *p < 0,05.

Overall, there is a substantial body of evidence about sex differences in many aspects of brain development and function and also behavior (reviews in Arnold and Chen, 2009; Ngun et al., 2011; Arnold, 2012; de Vries & Forger, 2015). Although such differences are mostly due to steroid hormone levels, recent data indicate that the sex-specific genetic architecture and expression of sex-linked genes also play a direct role on sexual dimorphism (e.g. de Vries et al., 2002; Gatewood et al., 2006; McPhie-Lalmansingh et al., 2008; Lentini et al., 2013; Corre et al., 2016; Arnold, 2019). The direct effect of the sex chromosome complement on behavior and brain has been mostly assessed using transgenic mouse strains, called the “Four core genotypes” model (see De Vries et al., 2002; Cox et al. 2014 for a description). But these strains show some limitations, including the fact that these mice are genetically manipulated, have poor fertility, and that the *Sry* gene is absent in XY females (Ngun et al., 2011). The sex determination system of *M. minutoides*, shaped by one million years of natural selection (Veyrunes et al., 2013), in which gonadal sex and chromosomal sex are partially dissociated, offers a unique framework to investigate the relationship between sex chromosomes, hormones and behaviors in a wild species. The hormonal profile of each female genotype obtained in this study confirmed that most of the sexually dimorphic behaviors of this species are driven by genetic factors rather than steroid hormone concentrations (Table 2). They also highlight the need to consider naturally occurring sex reversal with alternative phenotypes, as observed in the pygmy mouse or in the bearded dragon (Li et al., 2016), to better understand the large spectrum of behaviors under the direct influence of genetic factors and as such, to challenge the classical view of steroid hormone predominance in the induction of sex differences in non-gonadal organs and behavior.

**Table 2:** Hormonal profiles of the four sex genotypes. Comparison between hormonal profile expected according to behavioral traits they display, and hormonal profile observed.

|  | Hormonal profile expected |  |  |  |
| --- | --- | --- | --- | --- |
|  | Aggressive behavior | Anxiety-related behavior | Reproduction, maternal care | Hormonal profile observed |
| XX females | T ↘ | C ↗ | E ↘ | T →<br>C ↗<br>E → or ↘ |
| XX* females | T ↘ | C ↗ | E ↘ | T →<br>C ↗<br>E → |
| X*Y females | T ↗ | C ↘ | E ↗ | T →<br>C ↘<br>E → |
| XY males | T ↗↗ | C ↘ | NA | T ↗↗<br>C ↘ |

T = testosterone ; C = baseline corticosterone ; E estradiol ; ↗ = high concentration ; → = no difference between females ; ↘ = low concentration; double arrows indicate substantial difference.

## ACKNOWLEDGMENTS

We are grateful to RAM-CECEMA animal facility and Marie Challe for her help in maintaining the breeding colony. This study was supported by the ANR grant SEXREV (no. 18-CE02-0018-01).

